# Contrasting dissemination patterns of diploid cultivated cotton species *Gossypium herbaceum* L. and *G. arboreum* L. in Antiquity

**DOI:** 10.1101/2024.10.30.616484

**Authors:** Christopher R. Viot

**Author notes:** **Contacts** Christopher R. Viot. **Statements and Declarations**. Competing Interests: None declared. Permissions: None declared. Sources of Funding: None declared.

## Abstract

The textile use of cotton fibers from sister species *Gossypium arboreum* L. (Tree Cotton) and *G. herbaceum* L. (Levant Cotton) began, respectively, in Asia in the eighth millennium BP and in Africa in the third millennium BP and probably earlier. Archaeological data show that cotton cultivation spread in the Antiquity over most tropical and subtropical regions suitable for agriculture in Afroeurasia. The geographical expansions of the two Old World cotton species appear very contrasted, in terms of their chronologies, in terms of the areas conquered, and as for their speed of diffusion. *G. arboreum* spread over humid or sub-humid regions of southern, western and eastern Asia at *ca*. 0.8 to 1.7 km/yr, a rate close to that estimated for wheat from the Fertile Crescent to northwestern Europe during Antiquity, while *G. herbaceum* spread over arid to sub-arid regions of Africa and western and central Asia, much later and more than five times faster than *G. arboreum*. The very high speed of diffusion of *G. herbaceum* can be explained by the context and by the agricultural adaptations of this species but also leads to revisit the history of its domestication and beginning of cultivation.

## Introduction

The cotton species domesticated in Africa and Asia, *Gossypium herbaceum* L. (Levant Cotton) and *G. arboreum* L. (Tree Cotton), produce textile fibers whose use by humans began in the mid-Neolithic period. *G. herbaceum* and *G. arboreum* are sister species whose separation is estimated at *ca*. 0.7 Mya BP^1^ by molecular studies (Wendel 1989; Renny-Byfield et al. 2016; Huang et al. 2020) and which are neatly similar as for plant morphology and fiber characteristics but show clear genomic differentiations, including major chromosomic rearrangements (Gerstel 1953; Renny-Byfield et al.2016; Grover et al. 2022). They are the only diploids producing spinnable fibers within genus *Gossypium* (Fryxell 1979); the other two cultivated cotton species are tetraploids that were domesticated in the Americas and reached Afroeurasia recently through the post-Columbian exchanges. Textile fibers of the cultivated species of the genus *Gossypium* are botanically unicellular hairs, or trichomes, which grow from the seed coat and remain attached to the seed, unlike e.g. kapok (*Bombax* spp.). These long textile fibers or lint are different from the short, thick and strongly fixed on the seed, fibers, called fuzz, observable on the seeds of all species of the genus *Gossypium* (Hutchinson et al. 1947). Fuzz fibers are thought to be related to seed dispersal or germination control in relation to uncertain rain abundance, while the additional selective advantage linked to the much longer lint fibers remains undetermined, but has been proposed to be long-distance dispersal of seeds either by birds for their nests or by wind (Fryxell 1979; Hovav et al. 2008; Camacho et al. 2019).

The domestications of *G. herbaceum* and *G. arboreum* occurred independently in Northeast Africa and South Asia, respectively. Cotton fibers and threads dated to the first half of the eighth millennium BP were found in different funerary contexts at Mehrgarh in the Indus Valley and cotton cultivation seems to begin there in the middle of the seventh millennium BP (Costantini 1984; Moulherat et al.2002). In Africa, the oldest cotton fibers and seeds in an archaeological context were discovered at Afyeh, a Nubian site in the Nile Valley, and dated to the middle of the fifth millennium AD (Chowdhury and Buth 1971), without certainty however as to their date, origin, and human use; cotton cultivation is ascertained only in the second half of the third millennium BP in the region comprising northeastern Africa and northwestern Arabia (Bouchaud et al. 2018; Ryan et al. 2023). Cotton cultivation began to spread into Afroeurasia in the Antiquity, *G. arboreum* into southern, western and eastern Asia, and *G. herbaceum* into northeastern Africa and western and central Asia (Fuller 2008; Kulkarni et al. 2009; Bouchaud et al. 2018; Kelley 2018; Viot 2019). In spite of partly overlapping dissemination areas, and even in the cases of sympatric cultivation (Stanton et al. 1994; Kulkarni et al. 2009; Milon et al. 2023), *G. herbaceum* and *G. arboreum* kept nearly totally independent breeding trajectories as their genomic differentiation induces a strong reproductive isolation between them (Stephens 1949). As a consequence of human interest in the textile fiber they produce, the two Old World cottons *G. herbaceum* and *G. arboreum* eventually acquired a great importance in agricultural, artisanal and commercial activities and in long-distance trade and cultural networks of much of Afroeurasia until the beginning of the 20th century (Riello 2016; Bouchaud et al. 2018).

Archaeological data but also written evidence offer a record of the beginnings of cotton cultivation and its expansion over Afroeurasia in the Antiquity. The present work aims at precisely assessing and comparing the characteristics of the initial geographical disseminations during the Antiquity of the two domesticated diploid cotton species *G. herbaceum* and *G. arboreum*. It also intends to highlight which factors could have been determinant in how and where each species spread, such as geographic origins, agronomic adaptations and human context, and to elaborate from this comparison a better description and better hypotheses about the domestications and the first steps of cotton cultivation in Afroeurasia, in particular for species *G. herbaceum* for which our knowledge is much incomplete and the beginnings stay largely enigmatic (Bouchaud et al. 2018).

## Materials and Methods

The speed of the initial geographic expansion of cotton cultivation was assessed through the linear correlation between time and distance data of earliest proven cotton cultivation in sites over Afroeurasia for each diploid cultivated cotton species. Time data are the datations in years BP of the archaeological evidence and distances are the linear distances computed in the case of each species from its hypothetical domestication or initial cultivation center. The full list of early archaeological evidence about cotton use, their bibliographic sources, the locations, dates and distances, and the species involved, are given in table S1-A in Supplementary Material. The map of Figure 1 features the hypothetical geographic center wherefrom each cotton species began its dispersal, the hypothetical itineraries towards the different regions reached by cotton during its geographic expansion over Afroeurasia, and the as-the-crow-flies distances (in kilometers) between the main steps of these itineraries. The conclusions of the publications as regards the effective cotton cultivation in each site were strictly followed; local cultivation of cotton can generally be strongly suspected, if not demonstrated, when seeds, pollen and cotton textiles are abundant, or when texts clearly mention the cultivation (Ryan et al. 2023). Only the earliest occurrence of cotton cultivation in a given area was taken into consideration, so that some data corresponding to sites where the archaeological evidence of cotton cultivation was dated as much posterior (> one half-millennium) to surrounding sites weren’t included in the computations; the data effectively used are marked as such in Table S1-A, which also gives averages and standard deviations of time and distance for groupings of sites that were used for Figure 2.

**Fig 1.**
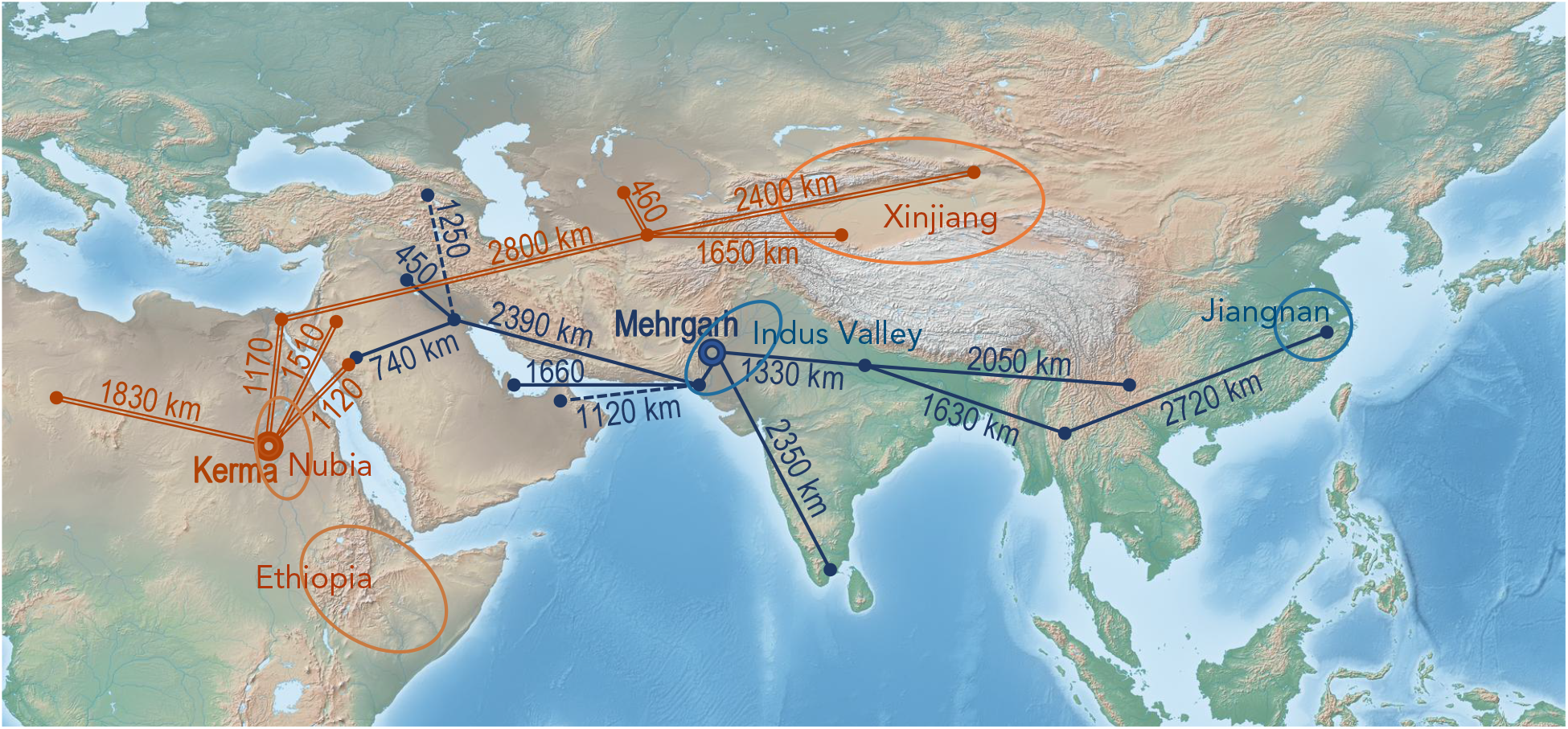
Distances along the hypothetical routes for the diffusion of cotton cultivation from the supposed initial places of cultivation or along networks for textile products exchange. Double lines for *Gossypium herbaceum*, solid lines for *G. arboreum* and dotted lines for cotton textiles exchange without local cultivation.

**Fig 2.**
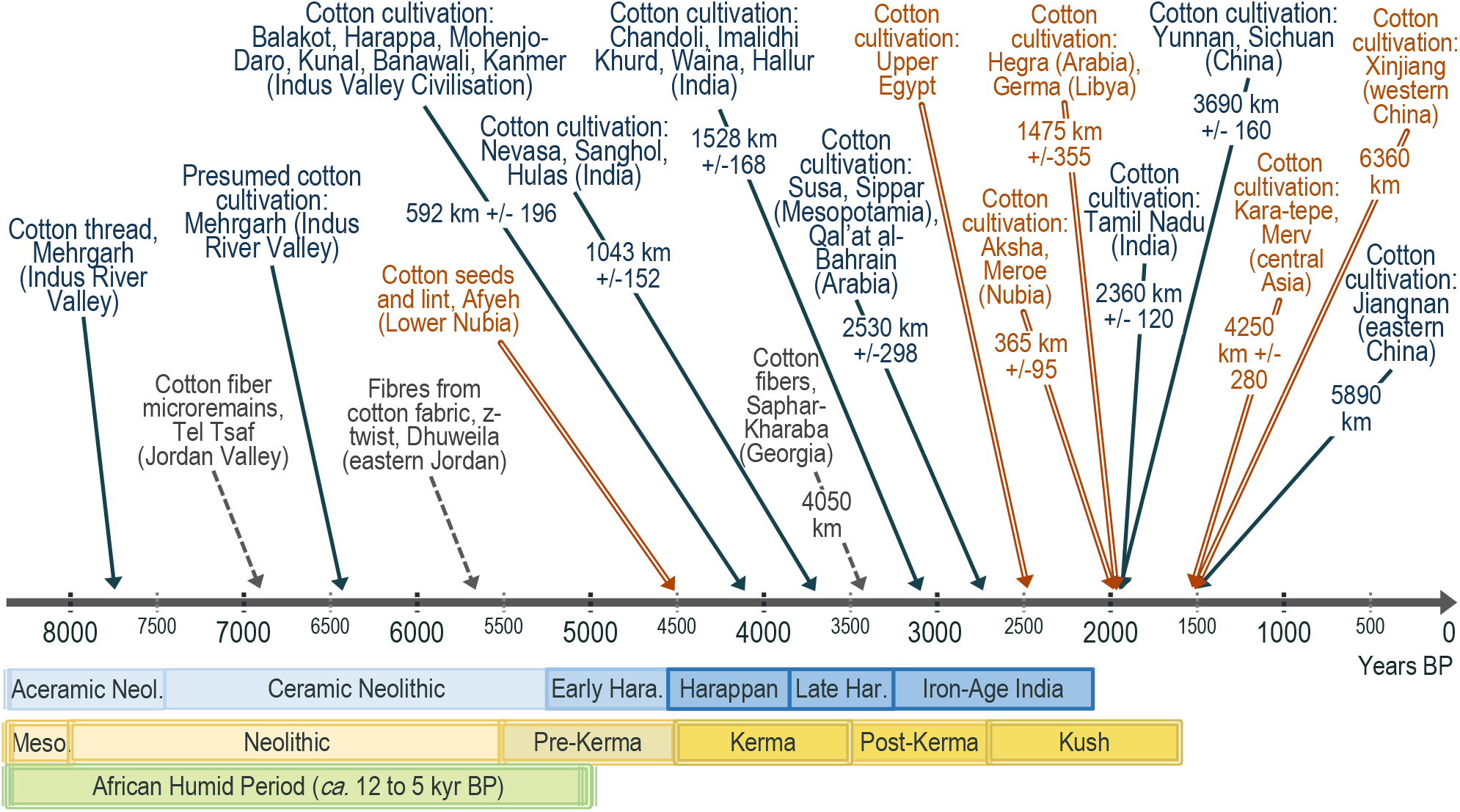
Schematic timeline of the expansion of the cultivation and textile use of cotton in the Old World in the Antiquity. Solid lines for *Gossypium arboreum*, double lines for *G. herbaceum* and dotted lines for archaeological remains without local cotton production. Time scale in years BP; average distances from the hypothetical domestication centers are computed from Mehgarh in Balochistan for *G. arboreum* and from Kerma in central Nubia for *G. herbaceum*. Sites^2^ were grouped according to the time when cotton cultivation was estimated to appear. The archaeological periods indicated below the time arrow are in blue for the Indus Valley and in yellow and green for Nubia. Precisions on data: see Table S1-A in Supplementary Material.

The itineraries that are featured were those that could seem logical or the simplest ones to link new cultivation regions to the initial cultivation centers of the considered cotton species, and the distance data is for each location the sum of the distances along the shortest itinerary. The Mandalay region of Myanmar was included as a step to reach the Jiangnan area in eastern China as the Chinese *G. arboreum* race *sinense* is considered derived from the race *burmanicum* of *G. arboreum* (Watt 1907). Distinguishing the two species in archaeological remains is challenging if not most often impossible on the basis on morphological data alone; fibers, textiles and charred seeds are the most frequent archaeological remains and characteristic features are commonly lacking (Milon et al. 2023).

Molecular genetics analyses, which permit the species to be directly determined with certainty (Palmer et al. 2012), are often difficult on archaeological remains concerning cotton and were until now seldom. Indirect proofs of the specific origin of ancient cotton remains such as the twist direction of a thread, the geographic origin and the epoch too often don’t allow an absolute certainty (Bouchaud et al. 2011; Kelley 2018).

Species assignations here was based on the consensus emerging in the publications from main specialists, as explained below. Data are all concordant about *Gossypium arboreum* being the cotton species initially cultivated in southern, eastern and southeastern Asia (Watt 1907; Chao and Chao 1977; Fuller 2008; Renny-Byfield et al. 2016). *G. herbaceum* was demonstrated as the species cultivated in the Antiquity in Nubia and its diffusion out of northeastern Africa seems to have begun only at the end of the third millennium BP (Palmer et al. 2012; Bouchaud et al. 2019; Ryan et al. 2023). The diffusion of cotton cultivation in southern, western and southeastern Asia until around the end of the third millennium BP should thus correspond to only *G. arboreum*. The cotton species early cultivated in the Persian Gulf and Mesopotamia is supposed to be *G. arboreum* because of the strong trade networks with India and linguistic studies showing that loanwords for cotton in Middle Eastern languages are of Indic origin, e.g. *pambakis, karpas, karpasos* (Muthukumaran 2016), and because the diffusion of *G. herbaceum* probably began centuries later. Nevertheless, which cotton was cultivated in Mesopotamia and in the Persian Gulf at different periods of the Antiquity still stays uncertain in the absence of confirmed species assignation (Bouchaud et al. 2018; Ryan et al. 2021), because of the lacking data about the history of *G. herbaceum* domestication and diffusion, and because in recent centuries *G. herbaceum* was the much dominant cotton in Mesopotamia and Persia (Kulkarni et al. 2009). *G. herbaceum*, first cultivated in Nubia, was demonstrated as the cotton species present in archaeological remains in Central Asia (Chao and Chao 1977; Cao et al. 2009); it is still unclear whether it was the cotton cultivated early in northern Arabia (Ryan et al. 2023), as was hypothesized by Kelley (2018). The cotton cultivated in Mada’In Salih (Hegra) at the end of the third millennium BP was not included in the computations as no hypothesis can reasonably be privileged about the species involved.

Distances were computed for *G. arboreum* from Mehrgarh, in province Balochistan of Pakistan. The archaeological site with the oldest remains of cotton textiles and oldest evidence of cotton cultivation in the Old World, and close to other archaeological sites which developed very early the use and cultivation of cotton, Mehrgarh most probably saw the domestication of *G. arboreum* (Costantini 1984; Moulherat et al. 2002; Fuller 2008). The region of Mehrgarh is also considered as one of the centers where agriculture first appeared during the Neolithic (Costantini 2008; Jarrige 2008). For *G. herbaceum*, Kerma was taken as the departure point for the distances to sites with new cultivation of this cotton species. Currently a small town in Sudan located on the east bank of the Nile, Kerma was the capital city of ancient Nubia during times when the country was flourishing and was more or less at its geographic center. Nubia is the region where the oldest traces of use and cultivation of *G. herbaceum* were discovered (Chowdhury and Buth 1970; Bouchaud et al. 2018; Yvanez and Wozniak 2019), and seems to have been wherefrom *G. herbaceum* initially disseminated towards other regions during the Roman domination over Egypt and the Levant (Kelley 2018). As for central and western Africa, too few data about cotton cultivation in the Antiquity were available for a correct knowledge of how cotton disseminated inside Sahelian and humid tropical Africa, as it is known it early did (Hutchinson 1949); data were similarly insufficient about cotton expansion into southeastern Asia.

Herodotus (430BCE) wrote about cotton in ‘Upper Egypt’ during Pharaoh Amassis’ reign (570-526 BCE, i.e. 2520-2474 BP); this was included in Figure 2 into the timeline of cotton development in northeastern Africa, as well as the cotton remains dated to 4550-4350 BP (Chowdhury and Buth 1971) found at Afyeh, southeast of Aswan and near to Qasr Ibrim, including seeds and fibers that were considered in the process of domestication by the authors. Although there are doubts about these data (Fuller 2015; Bouchaud et al. 2018, 2019), they are highly compatible with a past cultivation there of *G. herbaceum*, the cotton species that was domesticated in these regions of northeastern Africa. These data nevertheless are not involved into the computations of the diffusion.

## Results

### Timeline of the initial expansion of cotton cultivation in Afroeurasia

Figure 2 schematically summarizes the chronology of the development of cotton cultivation in Afroeurasia, according to archaeological data (sources: see table S1-A). Sites with evidence of cotton cultivation were grouped according to dates and geographical proximity, and averages computed for the dates of the archaeological evidence and for their distances from the hypothetical initial cultivation center for each species. As apparent in Figure 2, average distances are for each species increasing over time, as expected if the centers wherefrom the expansions were computed are close to the historical ones. We note that data were scarce for *G. herbaceum* (arrows with double line in Figure 2) and that its geographical expansion took place over a very short period, between approximately 2000 BP and 1500 BP, while the geographical expansion of *G. arboreum* (data with solid line arrows) began at around 4500 BP, reached the south of the Indian Peninsula at *ca*. 2000 BP and eastern China at *ca*. 1500 BP. Data with dotted line arrows relate to textile remains supposedly without local cultivation and derived from commercial or cultural exchanges.

The map of Figure 3 features the sites showing the earliest evidence of cotton cultivation in Asia and Africa, the centers where hypothetically their cultivations began and the probable pathways of the initial diffusion of the cultivation of the two Old Word cotton species according to archaeological and textual data (Herodotus 430BCE; Theophrastus 330BCE; Chowdhury and Buth 1971; Chao and Chao 1977; Moulherat et al. 2002; Tewari et al. 2006; Fuller 2008; Cao et al. 2009; Kulkarni et al. 2009; Palmer et al. 2012; Muthukumaran 2016; Kelley 2018; Ryan et al. 2023). Only very few dates correspond to cotton with confirmed species assignation (double lines for *G. herbaceum* and solid lines for *G. arboreum*). Ethiopia was featured here as some convincing proposal (Nicholson 1960) is that a wild *G. herbaceum* population in the southern lowlands of this country could have been domesticated there, giving the cultivated germplasm.

**Fig 3.**
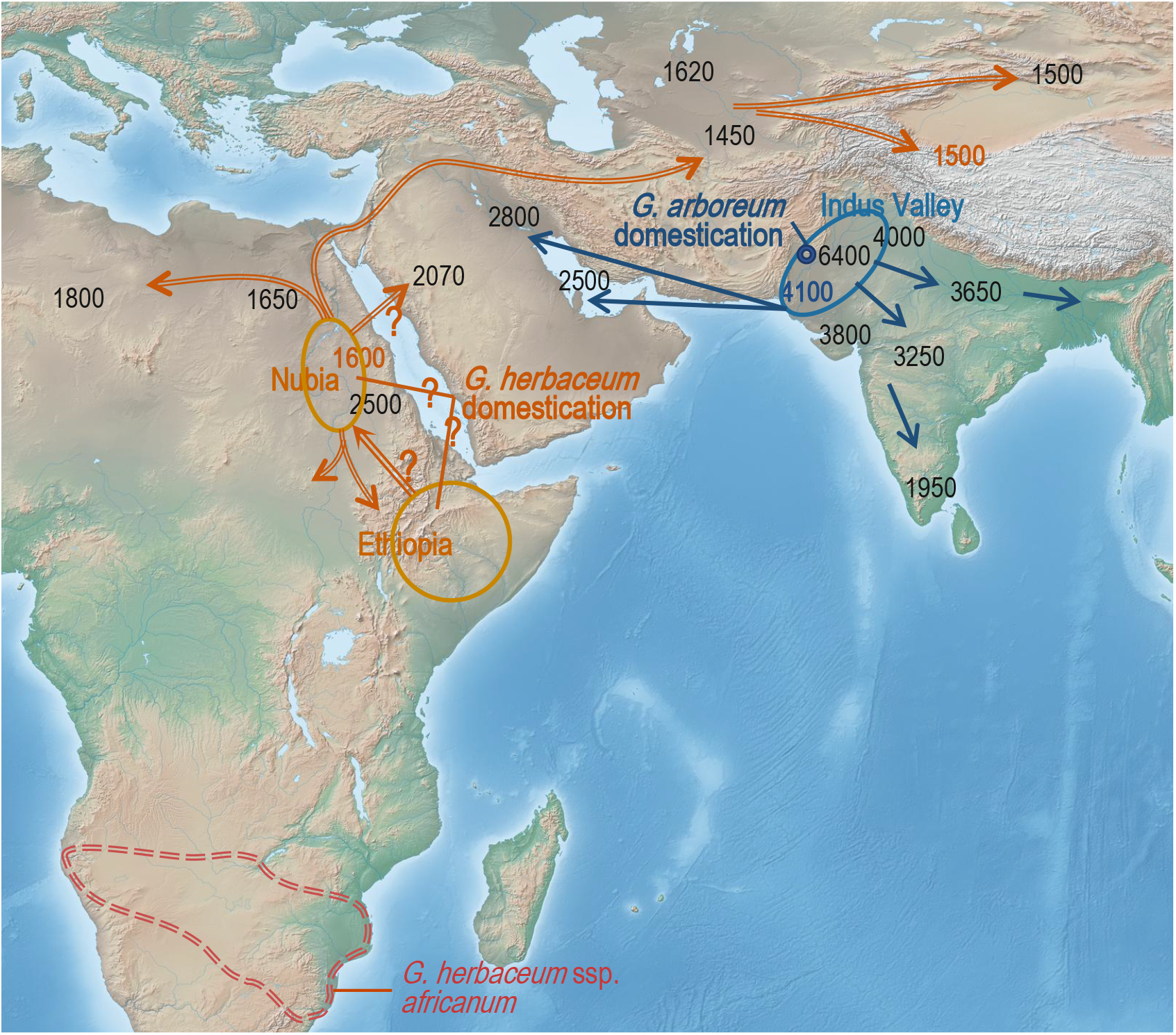
Earliest dates of cotton cultivation in Asia and Africa, hypothetical locations of the initial domestications of *Gossypium herbaceum* and *G. arboreum*, and probable diffusion pathways for each species. Double lines for *Gossypium herbaceum* and solid lines for *G. arboreum*. All dates are in years BP. The question marks indicate uncertainty about the place of domestication of *G. herbaceum*, and the possibility of its early diffusion towards northwestern Arabia. Sources: see text.

As appears in Figures 2 and 3, geography and time clearly separated the two Old World domesticated cottons before the end of the third millennium BP, *G. arboreum* having spread in Asia and *G. herbaceum* staying restricted into a small region of northeastern Africa. After the latter species began its spread, overlapping of the diffusion areas could occur. The cotton cultivated in the Persian Gulf and southern Mesopotamia in the third millennium BP most probably was *G. arboreum*, as explained above, and *G. herbaceum* necessarily crossed by Mesopotamia on its way towards Central Asia around the beginning of the second millennium BP. In northwestern Arabia, the cotton cultivations at around 2000 BP at Hegra (now Mada’In Salih, Medina province, Saudi Arabia) was either *G. arboreum* and then the Red Sea was the limit between this species and *G. herbaceum* cultivated in Nubia, or inversely the latter species was cultivated in Hegra and the Arabian Peninsula was then the contact region between the two diffusion areas. We still don’t know where and when the cultivation areas of cottons *G. herbaceum* and *G. arboreum* began to overlap, but reasonable candidates are thus Arabia and Mesopotamia around the beginning of the second millennium BP. The cotton in Hegra was in close geographic proximity to the cotton in Nubia at this time and these two regions nearly symmetrically positioned relative to the Red Sea are similarly arid such that the species *G. herbaceum* cultivated in Nubia might have been more adapted to the conditions in Hegra than *G. arboreum*; it was already proposed by Kelley (2018) that the early cotton cultivation in northwestern Arabia must have involved *G. herbaceum*. Cotton textiles from India, nevertheless, were traded in those times at Red Sea ports of Berenike and Myos Hormos (Bouchaud et al. 2011; Wild and Wild 2014), potentially bringing seeds of the species *G. arboreum* that was then grown in India, such that cultivation of the latter species in regions surrounding the Red Sea can’t be discarded.

### Dispersal rates of *G. herbaceum* and *G. arboreum* in Afroeurasia during Antiquity

The main parameters of the linear correlations between time and distance for the earliest occurrences of cotton cultivation in Afroeurasia are given in Table 1: correlation coefficient, slope, sample size and probability level. Slopes and correlation coefficients are negative because the time BP used for the computation is decreasing as dissemination distance increases from the initial cultivation sites. The slope corresponds to the average rate in km/yr - the linear speed in fact - of cotton geographic expansion. The linear correlations are significant at thresholds 0.025 or 0.005. For each species, in Table 1, the leftmost column gives the correlation parameters for all data while the columns just to the right are correlation parameters for subsets of the data: subdivisions in three epochs for *G. arboreum*, and computation only until Merv, without considering the step through to Xinjiang, for *G. herbaceum*.

**Table 1.** Linear correlation between distance and time of the initial spread of cotton cultivation in Afroeurasia, in Antiquity. Distances were computed in kilometers from hypothetical initial cultivation places, Mehrgarh for *G. arboreum*, and Nubia for *G. herbaceum* as explained in Material and Methods; time in years BP is the oldest archaeological record related to cotton cultivation in a region. The data used begin with the expansion of cotton cultivation to sites close to the hypothetical domestication place, not including thus the record of earliest cultivation in the domestication site itself.

|  | *G. arboreum*<br>4500 to<br>1500 BP | *G. arboreum*<br>4500 to<br>3000 BP | *G. arboreum*<br>3500 to<br>2500 BP | *G. arboreum*<br>3000 to<br>1500 BP | *G. herbaceum*<br>2000 to<br>1500 BP | *G. herbaceum*<br>Through to<br>Merv only |
| --- | --- | --- | --- | --- | --- | --- |
| Correlation coefficient | -0.874 | -0.873 | -0.852 | -0.767 | -0.877 | -0.870 |
| Slope (km/yr) | -1.3 | -0.8 | -1.0 | -1.7 | -8.4 | -6.7 |
| N | 22 | 13 | 15 | 13 | 7 | 6 |
| p | 0.005 | 0.005 | 0.005 | 0.005 | 0.025 | 0.025 |

### Initial *G. arboreum* dispersal over Asia

The earliest hints of cotton cultivation in the Old World, dated to the mid-7^th^ millennium BP in the Indus Valley, were evidenced at Mehrgarh, in Kacchi Plain (Gulati and Turner 1929; Moulherat et al. 2002). Clear signs of cultivation dated between 4500 to 3800 years BP, at the end of the Early Harappan phase, appear in archaeological sites belonging to the Indus Valley Civilization and distant from around 220 to 820 km from Mehrgarh (*ca*. 600 km on average), namely Balakot (on the Makran coast), Harappa, Mohenjo-Daro, Kunal, Banawali and Kanmer, as shown in Figure 2. In the following half-millennium, cotton cultivation is apparent in sites more distant (*ca*. 1000 km on average) and half a millennium later in sites even more distant (*ca*. 1500 km on average), all on the Indian subcontinent. Later, at *ca*. 2600 BP on average, cotton cultivation began to be present in Mesopotamia (Sippar), Persia (Susa) and the Persian Gulf (Qal’at al-Bahrain), on average at *ca*. 2500 km from Mehrgarh. At around 2000 BP cotton plants were present in most regions in the Indian subcontinent and also in southern China, in Yunnan and Sichuan provinces; around half a millennium later cotton was present in the Jiangnan region, east of Shanghaï. All data about cotton cultivation in these regions of Asia until then can be assigned to *G. arboreum*, as explained above in Material and Methods. This species seems to have been disseminated then only as a perennial crop (Watt 1907).

According to the linear regression of time and distance data, the overall speed of cotton cultivation expansion in southern, western and eastern Asia from *ca*. 4500 to 1500 BP was 1.3 km/yr (Table 1); the rate of expansion was slowest at the beginning, *ca*. 0.8 km/yr, from around 4500 to 3000 BP, and fastest in final steps, *ca*. 1.7 km/yr, from around 3000 to 1500 BP. These rates look comparable to those that could be observed for wheat (Cavalli-Sforza 1974; Kislev 1984; Crema et al. 2022). The rate of migration of wheat agriculture from the Near East to northern Europe was estimated an average 1.2 km/year (Kislev 1984); dividing the 3200 years of this expansion of wheat cultivation according to its major steps, 7600 BCE, 6750 BCE, 6000 BCE, 5100 BCE and 4400 BCE, Kislev (Kislev 1984) obtained rates that were 0.3, 0.8, 1.1 to 1.7 km/yr during each respective period; thus the increasing dissemination speed observed for *G. arboreum* cotton cultivation in our computations, from 0.8 km/yr to 1.7 km/yr, appears rather similar to that for wheat agriculture. The spread over Europe of wheat agriculture was dominantly linked to human migrations (Cavalli-Sforza 1974).

### Initial *G. herbaceum* dispersal over northeastern Africa and Asia

As shown by Figures 2 and 3, the initial expansion of *G. herbaceum* cultivation differs much from that of *G. arboreum*: beginning thousands of years later, its initial diffusion was completed in around half a millennium, reaching regions at distances similar to those reached by *G. arboreum*. It is only at around the change of era that the spread of *G. herbaceu*m out of Nubia begins, during the Roman domination over Egypt and the southern Levant (Bouchaud et al. 2018; Kelley 2018; Rohmer et al. 2022); in the following half-millennium it reached sites in southwestern Central Asia (Merv, Kara-Tepe) and in northwestern China (Hotan and Turpan in the Xinjiang).

Reaching the Xingjian in western China – *ca*. 5500 to 6500 km from Nubia in a rather straight line - at *ca*. 1500 BP corresponds to *ca*. 8.4 km/yr on average. Through to only Merv - around 4,000 km from Nubia and presumably the entry point to Central Asia - at *ca*. 1500 BP, average speed must have been at least 6.7 km/yr (see Table 1). The spread of *G. herbaceum* thus was around five times faster than that of *G. arboreum*. Uncertainties about the true dates when this species reached central Asia can be suspected as around two centuries separate the evidence for cotton cultivation in the rather close sites Merv and Kara-Tepe, and as cotton is evidenced nearly simultaneously in the Xinjiang, *ca*. 1500 to 2500 km further. It is much notable that the Xinjiang, in particular the regions around Turpan, is at latitudes excessively high (37 to 43°C) for cotton, with extremely cold winters. *Gossypium* cottons are intolerant to frost, and wild cottons in tropical regions have a short-day photoperiodism, inducing their flowering when days shorten and the fruiting in months that, out of the tropical zone, correspond to winter. Perennial habit and photoperiodism seem totally incompatible with climates such as in Xinjiang, such that it can only be supposed that annual, photoperiod-neutral *G. herbaceum* varieties were brought to Xinjiang. Watt (1907) already noted that the spread of *G. herbaceum* in inner Persia necessitated annualized forms.

### Long-distance commercial or cultural exchanges of cotton textiles

Cotton textile remains have been found in sites very far from the regions where cotton is known to have been cultivated at corresponding dates; these finds include 7150 to 6650-yr-BP fibers, partly dyed, at Tel Tsaf in the Jordan valley (Liu et al. 2022), 6400 to 4950-yr-BP fibers at Dhuweila in the Badiya region of eastern Jordan (Betts et al. 1994) and 3500 to 3400-yr-BP fibers in southern Georgia (Kvavadze et al. 2010). Cotton remains in human settlements at such early times in regions where cotton is not supposed to have grown wild or been cultivated cannot always be readily explained. For each case, the authors supposed that no local cotton cultivation could reasonably be hypothesized and, thus, long-distance commercial and cultural exchanges were to be involved. All these regions were included into diverse more or less overlapping trade and cultural networks extending around the 6^th^ to 4^th^ millennia BP over present-day northwest Iran, Mesopotamia, the southern Levant and the Early Transcaucasian Culture area, while already in the preceding millennia very active human migrations were expanding farming technologies to new regions (Yellin et al. 1996; Matessi 2023). These exchange networks and human migrations could have permitted cotton artefacts to reach sites much distant from production regions; cotton was highly priced, possibly for its dyeing properties and softness, so that it made much-valued gifts such as the ‘corselets’ described by Herodotus (430BCE). As exposed in paragraphs above, true cotton cultivation is supposed to begin in the mid-seventh millennium BP in Mehrgarh and in the mid-third millennium BP in Nubia, although the latter could have been the continuation of an earlier cultivation in Ethiopia (Nicholson 1960; Yvanez and Wozniak 2019). The cotton in southern Georgia at *ca*. 3500 BP fits perfectly with the hypothesis of commercial and cultural exchanges linking the Early Transcaucasian Culture area to Iran and Mesopotamia, as cotton cultivation was already well developed in the Indian subcontinent and could enter long-distance trading, the high latitude (*ca*. 42 °N) also being very unfavorable to cotton cultivation. The 6 to 5 kyr BP cotton fibers at Dhuweila in Jordan and the 7 to 6 kyr BP ones at Tel Tsaf in the Jordan valley also could come from the early phase of cotton production in the Indus Valley, but justifying this becomes really puzzling. A question could be whether a relationship exists between these last two archaeological cotton remains and the 4.5 kyr BP fibers and seeds at Afyeh in Nubia (Chowdhury and Buth 1971), because of their geographic proximity and although it involves an enormous timespan of three millennia.

## Discussion

The geographic expansions over Afroeurasia of cotton species *G. herbaceum* and *G. arboreum* in the Antiquity have been strikingly contrasted, as for the epochs, the durations and the rates in km/yr, and also as regards the regions covered. *G. arboreum* spread over a rather precisely known timespan, with a speed similar to that established for wheat during the Neolithic, over sub-humid or humid regions, and as, so far as is known, a perennial crop. *G. herbaceum* showed a comparatively extremely rapid dissemination which covered mainly arid regions and reached northern latitudes with agricultural conditions incompatible with cultivation of a perennial, photoperiodic cotton. The marked difference between the dispersals of the two Old World cotton species may supposedly find its explanation in their much-different historical and geographical contexts but also in the agricultural adaptions that differentiate the two species, as will be discussed below. The extremely brief timespan between the earliest certain cultivation of *G. herbaceum* in northeastern Africa and the culmination of its successful Afroeurasian dissemination which followed an apparent emergence in the middle of a hyper-arid region, after a nearly total absence from archaeological records before the end of the third millennium BP, also motivate a reexamination of the theories about its origins.

Our knowledge about the use of natural fibers and the cultivation of textile crops thousands of years ago can be strongly biased by the low rate of long-term preservation of organic matters such as fibers and seeds. The mostly cellulosic cotton fibers will be significantly degraded in a few weeks only, except in conditions of extreme dryness, coldness or low oxygen, or of protective mineralization (Reynaud et al. 2020; Margariti et al. 2023). Cotton seeds can also degrade rapidly or be eaten (by rodents, insects or birds) and their long-term preservation most commonly occurred as charred remains resulting from their carbonization (Milon et al. 2023). As a consequence, cotton fibers and seeds are rather infrequent in archaeological contexts and will most probably be detected for dates when cotton use and cultivation were becoming widespread rather than at early stages (Fuller 2008). The dates used here should thus be considered as minimal ages for cotton introduction in each region, as explained by Fuller (2008). The data about *G. arboreum* nevertheless depicted (Figure 2) a rather regular wave of advance of cotton over the Indian subcontinent and towards the Near East, which allows to think that a reliable representation of the phenomenon was obtained, even if some unknown number of years should be added to the observed ages. Another very critical point about reconstructing the history of Old World cottons is assessing which of *G. herbaceum* or *G. arboreum* was the species corresponding to the archaeological remains, and, as seen in Figure 3, few confirmed species assignations were available over the whole archaeological sites with cotton remains. Archaeogenomic studies are particularly lacking for Arabia and Mesopotamia as it becomes the contact zone between the diffusion areas of the two species in the second half of the third millennium BP.

### Cotton expansion and the Neolithic Revolution

The Neolithic Revolution, the shift from food gathering to food producing human communities, is generally estimated to have begun at *ca*. 12,500 years BP in the Fertile Crescent (Childe 1936; Fuller 2006); its eastward spread brought, *ca*. 3000 years later, Near Eastern domesticates such as barley, wheat and goats to the western Indian Subcontinent (Gangal et al. 2014), while local domestications included the zebu and jungle fowl, small millets and pulses, along with the *G. arboreum* cotton. The transition from nomadic hunter-gatherer societies to farming sedentary societies in the Indian subcontinent could be dated at between 9520 and 8150 BP in Bhirrana (Haryana state in India) and at between 8450 and 7450 BP at Mehrgarh, prehistoric sites both of the Indus basin (Gangal et al. 2014), and also had a very early beginning, at dates around the mid-9th millennium BP, in Lahuradewa in the Upper Gangetic Plain, around 850 km eastward of Bhirrana (Tewari et al. 2006). At *ca*. 4000 BP, agriculture was present in most studied sites of the Indian subcontinent, finally developing maybe only in the centuries before 3500 BP in Odisha, in eastern coastal India (Kingwell-Banham et al. 2018).

Cotton cultivation, thus, began in Mehrgarh around one to two millennia after subsistence agriculture began developing there, and extended around four millennia later than agriculture in those parts of the Indus Valley and the Gangetic plain where the cultivation of subsistence crops had a very early beginning; in other regions of the Indian subcontinent the spread of cotton cultivation and textile use seems to have followed by only a few centuries the development of general agriculture, which could have strongly influenced the speed of cotton cultivation progress. The spread of cotton cultivation over the Indus Valley, beginning at around 4500 BP, appears synchronous with the beginning of the Mature Phase (4550-2850 BP) of the Indus Valley Civilization (Figure 2). It is widely admitted that the establishment of agriculture was determinant for civilisation emergence (Westropp 1872) and the hypothesis of a strong link of civilisation development with textile production (Postrel 2020) could be much strengthed by the simultaneity observed here.

In northeastern Africa, there is evidence for crop cultivation in the oases west of the Nile in the eighth millennium BP (McDonald 2016), while cotton cultivation began in Nubia around five millennia later, in the second half of the third millennium BP (see Figure 2). Cotton wasn’t thus among the crops that directly followed the expansion of the Neolithic Revolution in these regions. In Egypt iself, cotton was not or nearly not cultivated until the Ptolemaic and Roman period, 305 BC-AD 395, while flax seems to have been the main textile crop (Melelli et al. 2021). An hypothesis suggested by Bouchaud (2020) about the absence of cotton in Egypt deals with a possible lack of workforce at the adequate moment for cotton harvest as their system of agricultural tasks was strictly dictated by the yearly flood of the Nile river and prioritized subsistence crops. From a biological point of view, it is notable that wet soils are totally inadequate for cotton plants, as they impact very negatively the root growth and function, so that cultivation in the flood-submersed zone could have appeared inadequate for cotton. Cotton cultivation higher on the river banks necessitated irrigation as all water for agriculture comes from the Nile river in this region where rains amount to a few mm/yr, and water-lifting means for irrigation, such as the saqiya, the noria, etc., began to appear by the 4th century BCE in Egypt, in the oases west of the Nile and in Arabia (Bouchaud 2020).

### The ingredients of *Gossypium herbaceum* rapid dissemination

The spread of *G. herbaceum* started at the end of the third millennium BP when long-distance commercial communication networks were already linking the southern Levant to northern Mesopotamia through to Central Asia, including particularly the Silk Routes that began in the last two centuries of the third millennium BP (Church et al. 2018), and must have facilitated the diffusion of this cotton species. The Silk Routes network corresponds rather precisely to the diffusion area of the African cotton in the Antiquity. The late diffusion of this cotton also occurred over regions were agricultural techniques were already mastered, possibly facilitating the adoption and geographic progress of the new crop; on the contrary, the diffusion of *G. arboreum* in South Asia seems to have partly occurred not long after agriculture itself, with a rather slow progression. The African cotton species clearly shows a better adaption than *G. arboreum* to arid or sub-arid continental environments (Watt 1907; Brite and Marston 2013), that could have made its diffusion partly by going from one oasis to the next, thus over rather long distances at each step. Diverse factors may thus have combined to strongly accelerate the spread of *G. herbaceum*.

### Spread of *G. herbaceum*, archaeological remains and hypotheses about its domestication

As shown in Figure 2, around one thousand years passed between the first documented textile use of cotton at Mehrgarh and the first hints of cotton cultivation in this same location, and then a *ca*. two-millennia more until the beginning of the expansion of cotton cultivation in the Indus Valley (Costantini 1984; Moulherat et al. 2002); similar long delays were observed for the tetraploid cottons that were domesticated more or less simultaneously, but of course totally independently, in tropical America (Viot and Wendel 2023). These long delays now appear as a common feature of crop domestication (Fuller 2010; Meyer et al. 2012). A rather recent concept is that there was a phase of proto-agriculture or proto-farming during which humans harvested wild plants they cared for, a system named ‘plant husbandry’, before true cultivation developed (Crawford 2011; Beldados et al. 2023). In the case of cotton, the plant husbandry phase would have permitted the evolution of the incipient crop towards forms that can be more easily grown and disseminated and also permitted the mastering of technologies for the textile use of the short cotton fiber.

The spread of *G. herbaceum* cultivation began a short time after the first certain signs of its cultivation in Nubia, and from then its cultivation extended in a few centuries into particularly difficult agricultural environments such as those of Central Asia. Questions are inescapable about this spread of *G. herbaceum*. First, how much did the wild *G. herbaceum* have to evolve before its spread and its easy cultivation out of tropical regions? Wild *Gossypium* cottons have characteristics that make their cultivation very difficult, or even impossible in some environments where *G. herbaceum* was quickly introduced, as explained above in §’Initial *G. herbaceum* dispersal over northeastern Africa and Asia’. In particular, the intolerance to frost will prohibit the survival of perennial forms in regions with cold winters and the short-day photoperiodism will impede fiber production in regions with a short hot season, as cotton stops growing and developing at temperatures lower than around 20°C (Reddy et al. 1991). For all domesticated cottons, domestication had to eliminate the physiologically-regulated seed dormancy and the hard seed coat, and also, for some regions the short-day photoperiodism, while other important traits such as seed and boll size, lint length, quantity (fiber turnout) and whiteness, and productive morphology, are less critical. Second, how long must be this domestication phase? From the observations on the other cotton species, we would estimate that many centuries are necessary, this of course in the case where all conditions are identical. At least, it seems that the domestication cannot have taken place in the region where.

The contradiction between what is known about the beginning of *G. herbaceum* cultivation and the usually long delays before a wild cotton species becomes easily cultivated and spread into different agricultural environments might push to search how and where the domestication phase could have taken place. The Nubian desert and close surrounding regions were over the last millennia before cultivation began rather unfavorable to a population of wild cotton plants. The wild subspecies of *G. herbaceum* ssp. *africanum* in southern Africa (see Figure 3), although considered the reasonable model for the ancestor of the cultivated germplasm (Hutchinson et al. 1947), cannot be the direct ancestor of the domesticated *G. herbaceum* germplasm, and the most convincing hypothesis was recently of a wild *G. herbaceum* population in Ethiopia that went domesticated (Nicholson 1960; Yvanez and Wozniak 2019). Crop cultivation appeared in Ethiopia at around 2500 BP (Harrower et al. 2010); cotton is in fact considered there as an old traditional crop (Nicholson 1960), the beginning of which nevertheless stays undetermined and it was cultivated only as a perennial crop until recently. An alternative hypothesis, best coherent with the archaeological finds, including that of very early cotton fibers in the Jordan valley, and with the observed timeline of cotton cultivation and diffusion, can be proposed. During the African Humid Period (AHP), from *ca*. 14,500 to 5000 years ago, yearly rainfall in what is now the hyper-arid Sahara was 300–920 mm (Ritchie et al. 1985; Ritchie and Haynes 1987), and these sub-humid conditions sustained a savannah-like environment with thriving human populations (Ghoneim et al. 2024), while the Nile Valley itself was on the contrary unfavorable to human settlements because of the swampy environment and frequent floods (Williams et al. 2010; Zaki et al. 2021). The Sahelian environment west of the Nile River valley during the AHP was probably suitable for populations of wild *Gossypium herbaceum*, as the Sahel now hosts the very close wild species *G. anomalum*. When the Sahara progressively became too dry, at *ca*. 6 kyr to 5 kyr BP, humans migrated from there, eastwards to the Nile banks, or to the South and West and to the Mediterranean (Kuper and Kröpelin 2006). A hypothetical *G. herbaceum* cotton growing wild and used by humans in the humid Sahara of the AHP, later brought to the Nile valley when the Sahara was drying up, could have been cultivated at a small scale until its evolution into a manageable crop. The *ca*. 7 to 6-kyr-BP cotton remains in Tel Tsaf in the Jordan valley (Liu et al. 2022) and the *ca*. 6 to 5-kyr-BP fibers found at Dhuweila in eastern Jordan (Betts et al. 1994) were contemporaneous with the last millennia of the African Humid Period; and the *ca*. 4500 BP seeds and fibers in Afyeh (Chowdhury and Buth 1971) become compatible with the populations retreating to the Nile River banks when the AHP was ending. The earliest cotton remains in this region and the timeline of *G. herbaceum* diffusion would be fairly better explained by this hypothesis of a proto-agriculture of wild cotton plants in the AHP Sahara than by exchanges with India at extremely early times and a very recent domestication with an excessively short period for the evolution of *G. herbaceum* into an agriculturally manageable plant. Whenever was then the 12-to-5-kyr-BP Humid Sahara the cradle for African cotton domestication, remains related to human activities and to flora and fauna in regions surrounding the Nile valley during the AHP would be now with much difficulty observable, being either destroyed by the desertification and sand dune activity or buried beneath sand sheets (Ghoneim et al. 2024).

## Conclusions

In this study, we assessed the speed of cotton cultivation spread into Afroeurasia in the Antiquity for each of the two domesticated Old World cotton species. It permitted to better understand the context, characteristics and differences of their disseminations and to establish how the epochs of the disseminations and the agricultural adaptations of each species were key factors. The markedly particular dissemination pattern shown by *G. herbaceum* permits to propose an alternative hypothesis for its domestication history, explaining the much missing data about its first steps and better compatible with the very early archaeological cotton remains in this region. Further progress about cotton in the Antiquity in Afroeurasia strongly needs DNA-based species assignation of archaeological remains.

## Acknowledgements

Background maps of Figures 1 and 3 are from Natural Earth project (https://www.naturalearthdata.com)

## Supplementary data

### S1 – Archaeological data on initial dispersal of domesticated Old World cottons

**Table S1-A.**
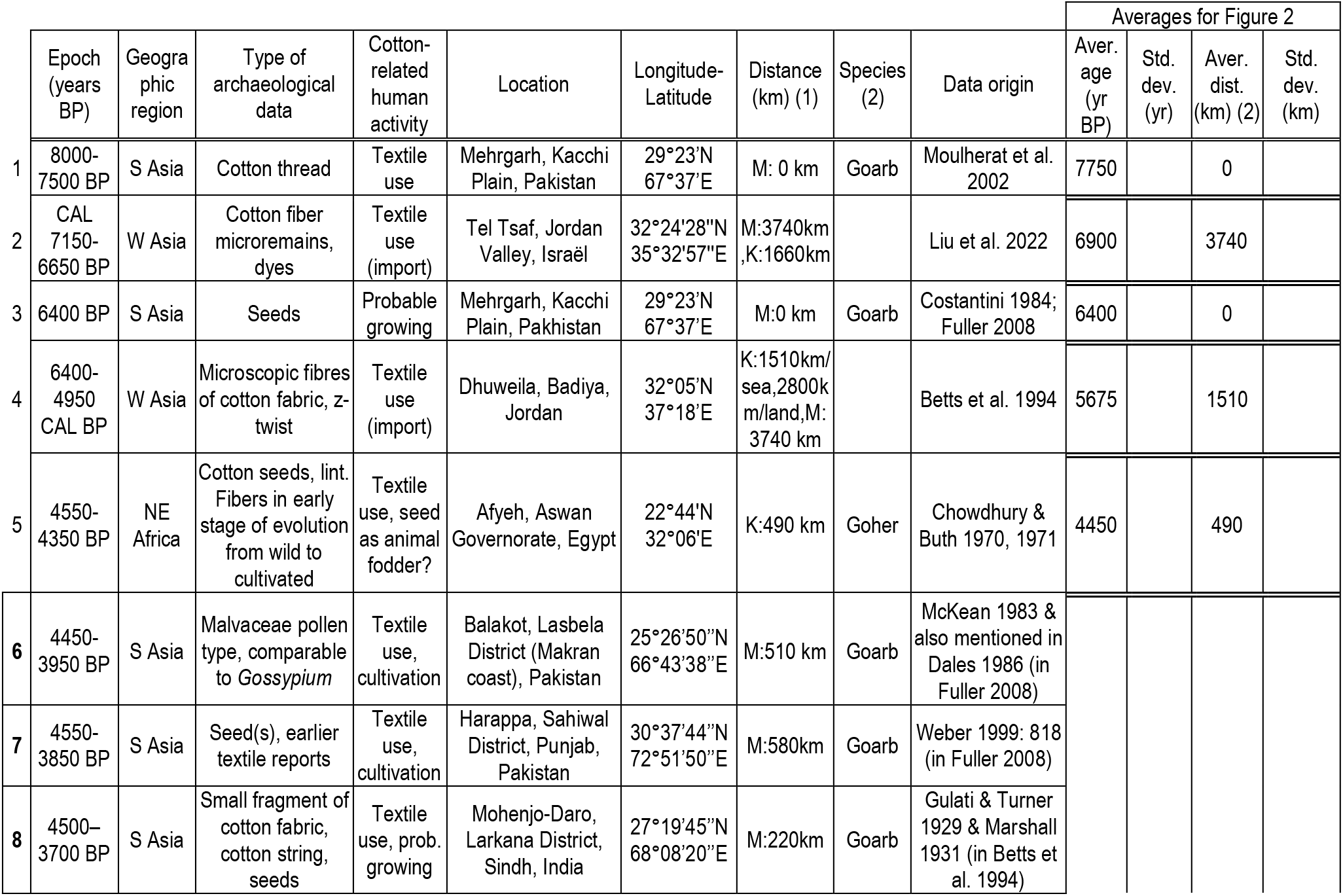
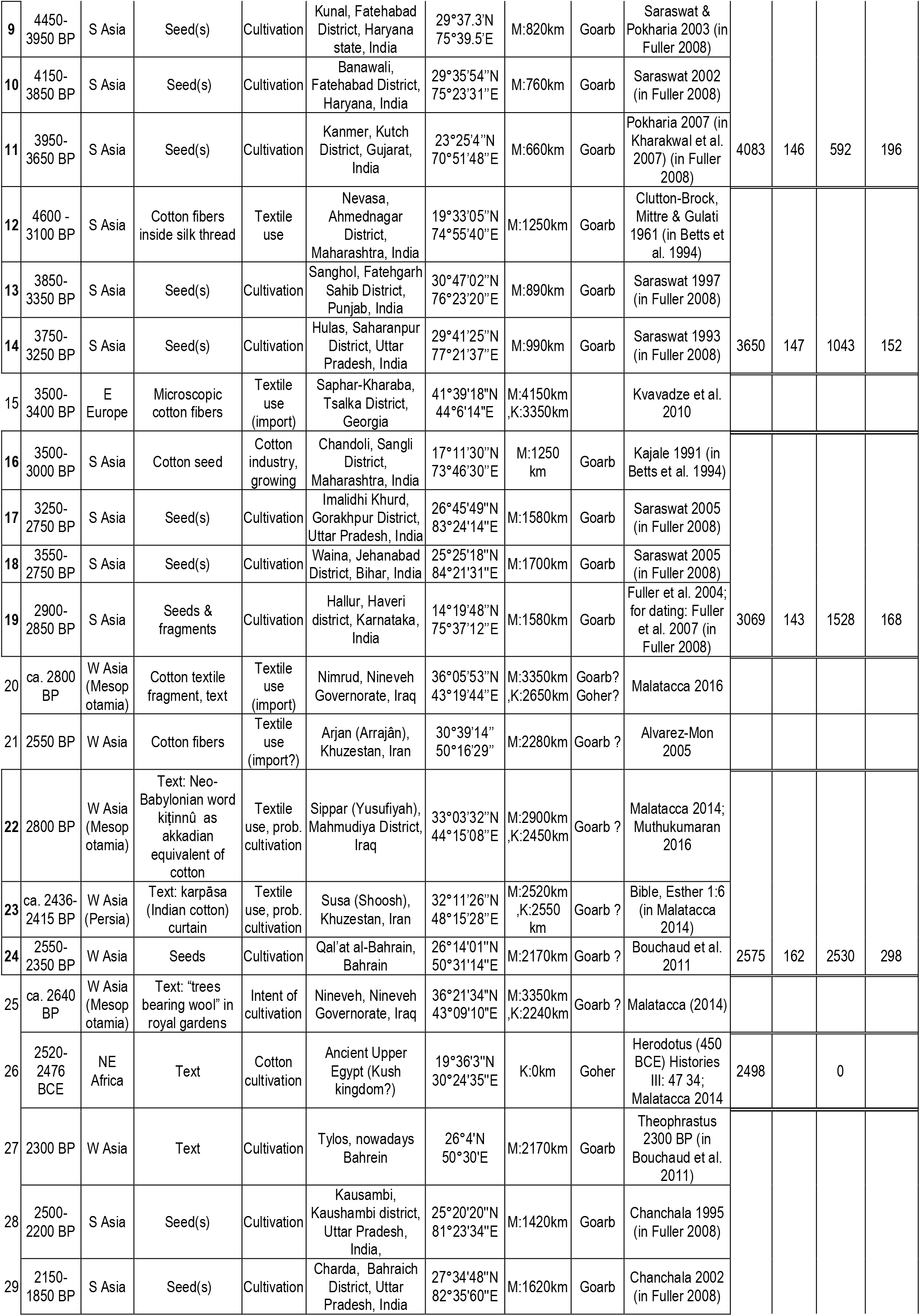
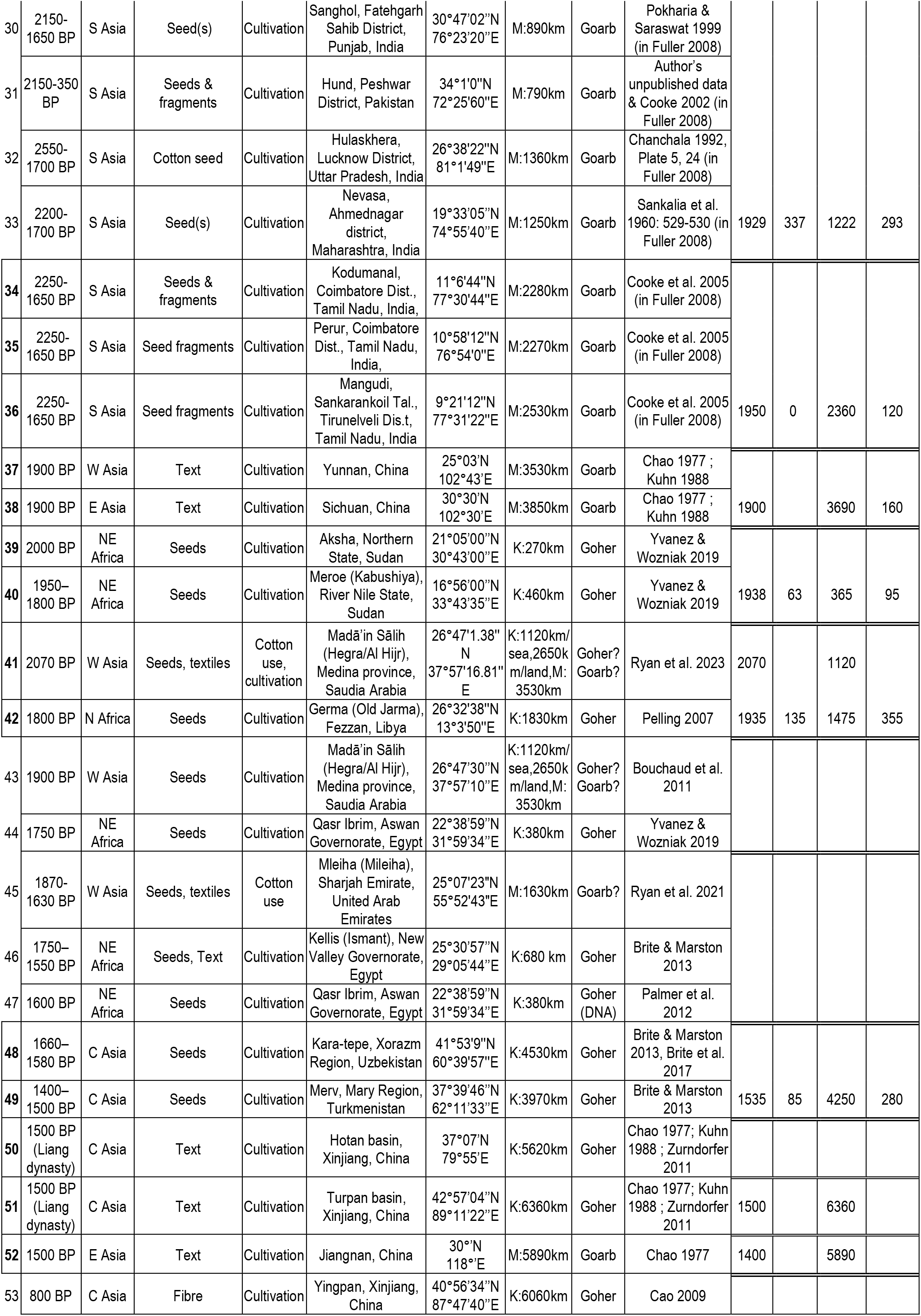
List of archaeological sites, cotton-related evidence, precise locations, times and distances. Table S1-A list the main oldest archaeological evidence of cotton cultivation and use (textile or other) in the Old World. These data were used to compute the characteristics of the initial geographic spread of the two diploid cultivated cottons. Data included in the computations are indicated by the line number in bold character and framed. In particular, data for the time-distance correlation begin with the first proven cultivation distant from the hypothetical center of the dispersals, and for a given geographic zone, only the earliest date with attested cotton cultivation was taken into consideration in the computations. Dates are in years BP and distances in metric kilometers; distances are from Mehrgarh in Pakistan (M: xx km) for *G. arboreum* and from Nubia (computed as from Kerma, K: xx km) for *G. herbaceum*, computed using https://www.google.com/maps. Averages and standard deviations of time BP and distances are indicated for groupings of the sites featured in Figure 2. Abbreviations: Aver.= Average; yr BP = years Before Present; Std. dev. = Standard deviation; dist.= distance; km: metric kilometers; yr: year; M: xx km = distance from Mehrgarh; K: xx km = distance from Kerma in Nubia; Goarb = *G. arboreum*, Goher = *G. herbaceum* (when not proven by studies, the species assignation is the most probable in accordance with authoritative works and scientific consensus, as explained in §Materials and Methods.)

## Footnotes

1 BP: Before Present i.e. calendar years before 1950; Mya: one million years ago, kya: one thousand years ago; years BCE, CE: years Before Common Era, of Common Era, corresponding to year notations for the Gregorian calendar. Dates are given in CAL years for radiocarbon-based datations.

2 Geographic precisions on cited sites (in alphabetical order): Aksha, Nubia; Balakot (Makran coast), Balochistan, Pakistan; Banawali, Haryana, India; Chandoli, Maharashtra, India; Charda, Punjab, India; Germa (Old Jarma), Fezzan, Libyan Sahara; Hallur, Karnataka, India; Harappa, Punjab, Pakistan; Hegra (Mada’in Salih), Saudia Arabia; Hulas, Uttar Pradesh, India; Imladhi Khurd, Uttar Pradesh, India; Jiangnan, eastern China; Kaushambi, Uttar Pradesh, India; Kanmer, Gujarat, India; Kara-tepe (Khorezm), Karakalpakstan, Uzbekistan; Kunal, Haryana, India; Mergarh, Balochistan, Pakistan; Meroe, Nubia; Merv, Turkmenistan; Nevasa, Maharashtra, India; Nineveh, Iraq; Qal’at al-Bahrain, Bahrein; Sanghol, Punjab, India; Saphar-Kharaba, Georgia; Sichuan, southern China; Sippar, Babylonia, Iraq; Waina, Bihar, India; Xinjiang (Turpan, Hotan,Yingpan), western China; Yunnan, southern China.

